# Changing rounds into squares or combining stripes? In-depth analysis of *Fritillaria* flowers and the color-changing pattern of *Sarcophaga* flies sheds light on the diversity and morphogenesis of checkerboard patterns in Eukaryotes

**DOI:** 10.1101/2024.02.07.579346

**Authors:** Pierre Galipot, Julie Zalko, Laetitia Carrive, Romain Garrouste

## Abstract

Important in many human artistic cultures, checkerboard patterns are rare in nature like many motifs based on squared geometry. Nevertheless, they are expected to be very detectable by the visual systems due to their periodic geometry and contrasted two-tone coloring, therefore potential specific biological functions are suspected. Here, thanks to a biological survey, we draw the first diversity landscape of eukaryotic species bearing checkerboard patterns, confirming their rarity but also their presence in extremely diverse clades. Then, we selected two genera, *Sarcophaga* flies and *Fritillaria* flowers, to perform in-depth pattern analyses allowing us to make strong hypotheses on the mechanisms producing these very peculiar patterns, as no morphogenetic process was known to generate checkerboards. Although they share a similar geometry, these two genera appear to produce checkerboards through very different ways, showing a convergence of shape but not of processes. Whereas the *Fritillaria* analysis points to a geometric constraining of a Turing-like pattern by the parallel network of veins, that of *Sarcophaga* suggest the reuse of developmental boundaries and right-left symmetry, together by the combination of vertical and horizontal stripes. Furthermore, we present the first description to our knowledge of the striking color-changing nature of *Sarcophaga* checkerboards, whose light and dark squares can exchange their color depending on the angles of lighting and observation thanks to the planar polarity of the cuticular hairs, the setae.

Together, this shows the extent of the processes selected during evolution to generate complex forms and colors, and confirms the importance of studying morphogenesis with in-depth pattern analyses and through species diversity. Finally, by enabling strong hypotheses to be made about the morphogenesis of these patterns, it paves the way for the molecular identification of the morphogenetic processes at work.

## Introduction

Contrary to hexagons and circles, square patterns are much more uncommon in nature. Many studies have proven the difficulty to produce square geometry and the instability of square lattices (1,2) (i.e. when each unit has 4 closest neighbors, and therefore often exhibit square shape). Nevertheless, cells exhibiting square lattice organizing and even checkerboard patterns have been described in some eukaryotic tissues, like the mouse auditory epithelium (3), the oviduct epithelium of the Japanese quail (4) or brown algae early developmental tissues (5). Checkerboard color patterns are the combination of a square lattice and two other characteristics (Fig 1B): square-shaped motifs and an alternation of two colors or shades (in case of coloration based of shades of grey). They are expected to be very detectable by the visual systems due to their periodic geometry and contrasted two-tone coloring (6–8). Two possibilities then coexist, either these pattern characteristics are directly produced by the same processus (in the same way that dots and their hexagonal arrangement are produced by the same process during Turing-like patterning (9)), or they are produced by two or more separated processes, simultaneously or not, which then can add up and/or interact to form checkerboards.

**Fig 1.**
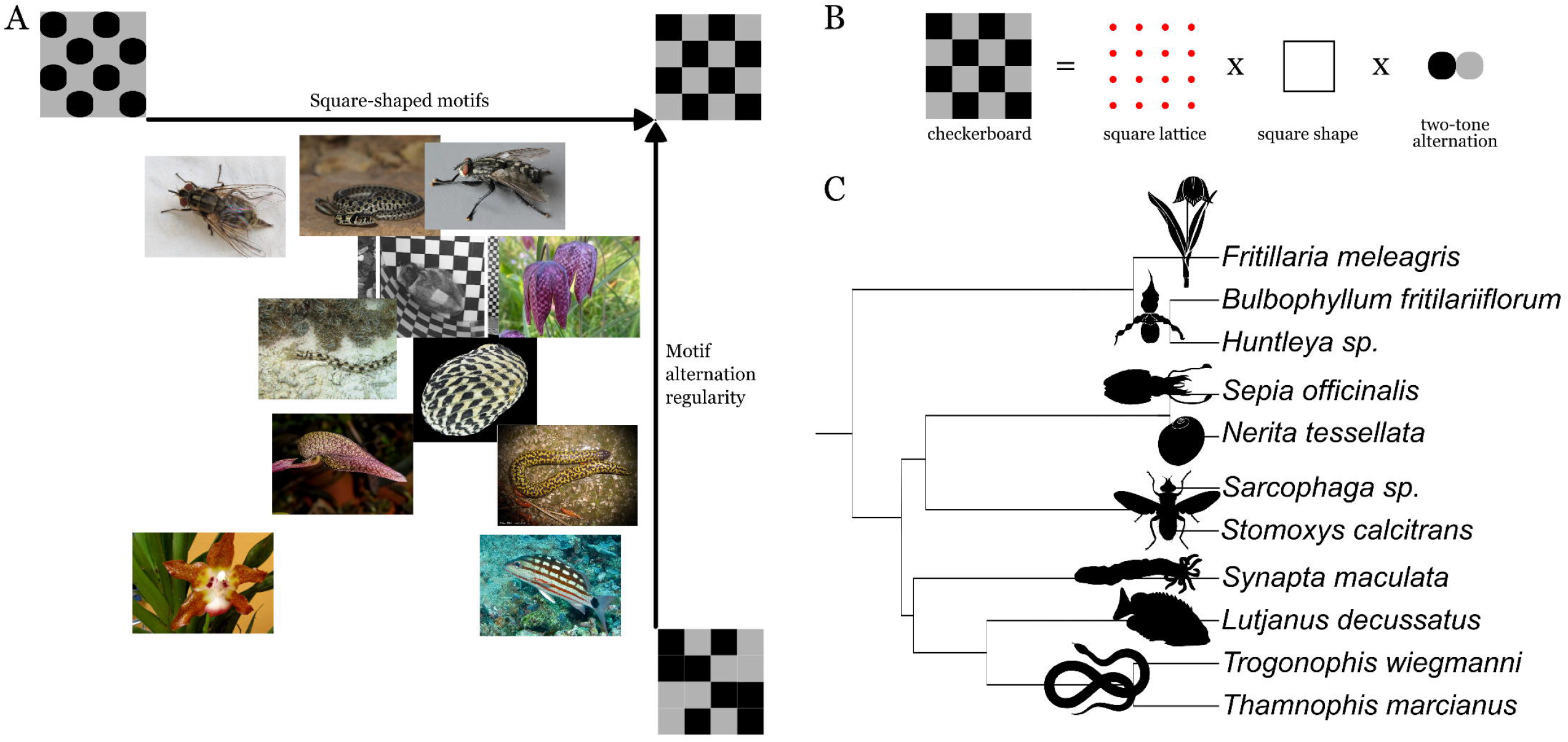
Diversity of Eukaryotes species bearing checkerboard-like patterns. (A) Semi-quantitative diagram of eukaryotic species bearing checkerboard-like patterns, according to the deviation of their pattern from the perfect mathematical pattern. (B) The three necessary and sufficient characteristics of a checkerboard pattern (C) Phylogeny of the selected species

If some species are quite famously known for their checkerboard-like color patterns like the *Fritillaria* flowers (10) (Liliaceae) which bear them on their tepals, few species have been formerly described for those peculiar shapes, and no work has already presented them combined in a single study.

We therefore chose to start with a description of as many biological checkerboard patterns as possible, using multiple data sources, taking care to cover the maximum of eukaryotic biodiversity. While the shortcut to molecular biology is tempting to highlight morphogenetic mechanisms, we advocate to start with an approach based on exploring diversity, known as the SE (せ) method (11), which then allows us to select the most suitable species for pattern analysis and functional biology. Due to the highly integrative nature of patterns and forms, morphogenesis is a field that rarely allows for the effectiveness of a bottom-up approach, or one focused on a single scale (12).

After the species survey, a large range of potential species to study was available and we decided to choose and compare an animal and an angiosperm species (and their respective genera): *Sarcophaga* flies and *Fritillaria* flowers. They presented the patterns closest to a perfect checkerboard motif (Fig 1A), and fresh and collection material were accessible to analysis. Microscopy observations, in-depth pattern analyses, simulations and species comparison allowed us to propose two very distinct mechanisms of formation, thanks to a cluster of clues. While the checkerboard pattern of fritillaries is formally described by multiple sources and even incorporated into the genus name (*Fritillaria* comes from “chessboard or dice box” in Latin (13)), *Sarcophaga* checkerboards are rarely mentioned in the literature. Furthermore, surprisingly, their color-changing nature has never been mentioned to our knowledge, perhaps due to the need to change the angle of lighting in order to observe it.

## Results

### Checkerboard color patterns are rare but present in many diverse taxa of Eukaryotes, notably plant and animal species

To perform a survey of species bearing checkerboard patterns, we first had to define them properly. We chose to consider that three characteristics are necessary and sufficient: i) motifs organized in square lattices (i.e. when each unit has 4 closest neighbors), ii) square-shaped motifs and iii) an alternation of two colors or shades (Fig 1B). Perfect checkerboard patterns are never reached by biological patterns; therefore, we chose to work with a tolerance to variation as biological patterns could not match perfect mathematical ones. Consequently, we include species with slightly diverging characteristics, like unperfect squares or partly irregular color alternation (Fig 1A). Since no exhaustive survey was possible, we chose to use keywords to find the species in the literature and biodiversity databases (like ‘checkerboard’, ‘checkered’, ‘check’, ‘square’ and ‘*Fritillaria*’, the Latin word for ‘dice cornet’, an object exhibiting a checkerboard pattern and then used in species names to pinpoint the presence of this particular pattern, or one close to it), together with surveys also sent to expert researchers in various taxonomic groups (typically working in natural history museums).

Thanks to this, we described several species and genera, summarized in Fig 1, and qualitatively represented in a diagram thanks to the more or less perfectness of their characteristics. Both Angiosperms and animals were found to exhibit checkerboard-like patterns, but we were not able to find any species belonging to Fungi. Very diverse animal species (vertebrates, insects, echinoderms, cephalopods, gastropods) were found, but few plant species seem to bear such patterns. The two most perfect checkerboard-like patterns were found in *Sarcophaga* flies genus, exhibited on their upper abdomen, and *Fritillaria* genus on their tepals.

We then chose these two groups to explore checkerboard patterns formation.

### Color patterns of the 164 recognized species of Fritillaria exhibit a large diversity of hues and shapes, including checkerboard whose morphogenesis is compatible with a Turing-like mechanism constrained by a parallel nervure frame

Before focusing our study on *Fritillaria meleagris*, we decided to characterize the flower color patterns of the 164 recognized species according to the Kew Botanical Garden POWO website (14). By harmonizing the descriptive terms for colors and using various sources of photographs, we described fourteen different colors of backgrounds and motifs (brown, purplish-brown, purple, pink, red, reddish-brown, green, yellowish-green, greenish-yellow, yellowish-green, yellow, orange, blue and white, see Fig 2A). Certain color combinations of pattern and background were more common than others, and in an intuitive way, strong contrasts in terms of brightness and complementary colors seem to have been favored during evolution (such as dark red-brown patterns on a light yellowish-green background, see Table 1).

**Fig 2.**
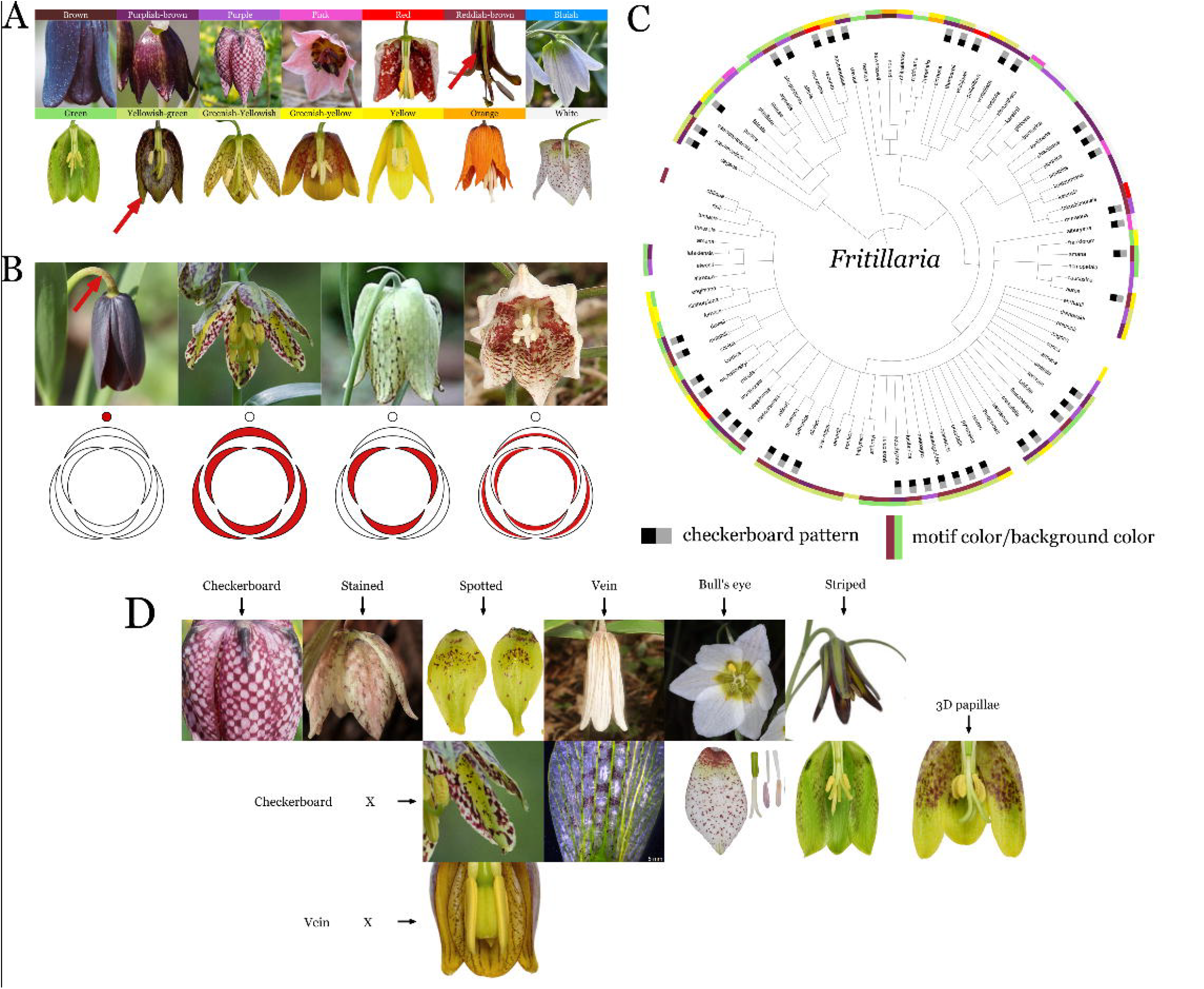
Diversity of colors, shapes and tissue location of Fritillaria color patterns. (A) The 14 harmonized hues of *Fritillaria* color patterns and their reference pictures (B) Four variations of pattern location, from left to right: peduncle, both sides of petal and tepal, both sides of petal only, internal sides of petal and tepal (C) Presence of a checkerboard pattern and motif and background colors mapped on *Fritillaria* phylogeny (D) The seven pattern categories and their possible combination with checkerboard patterns

**Table 1.**
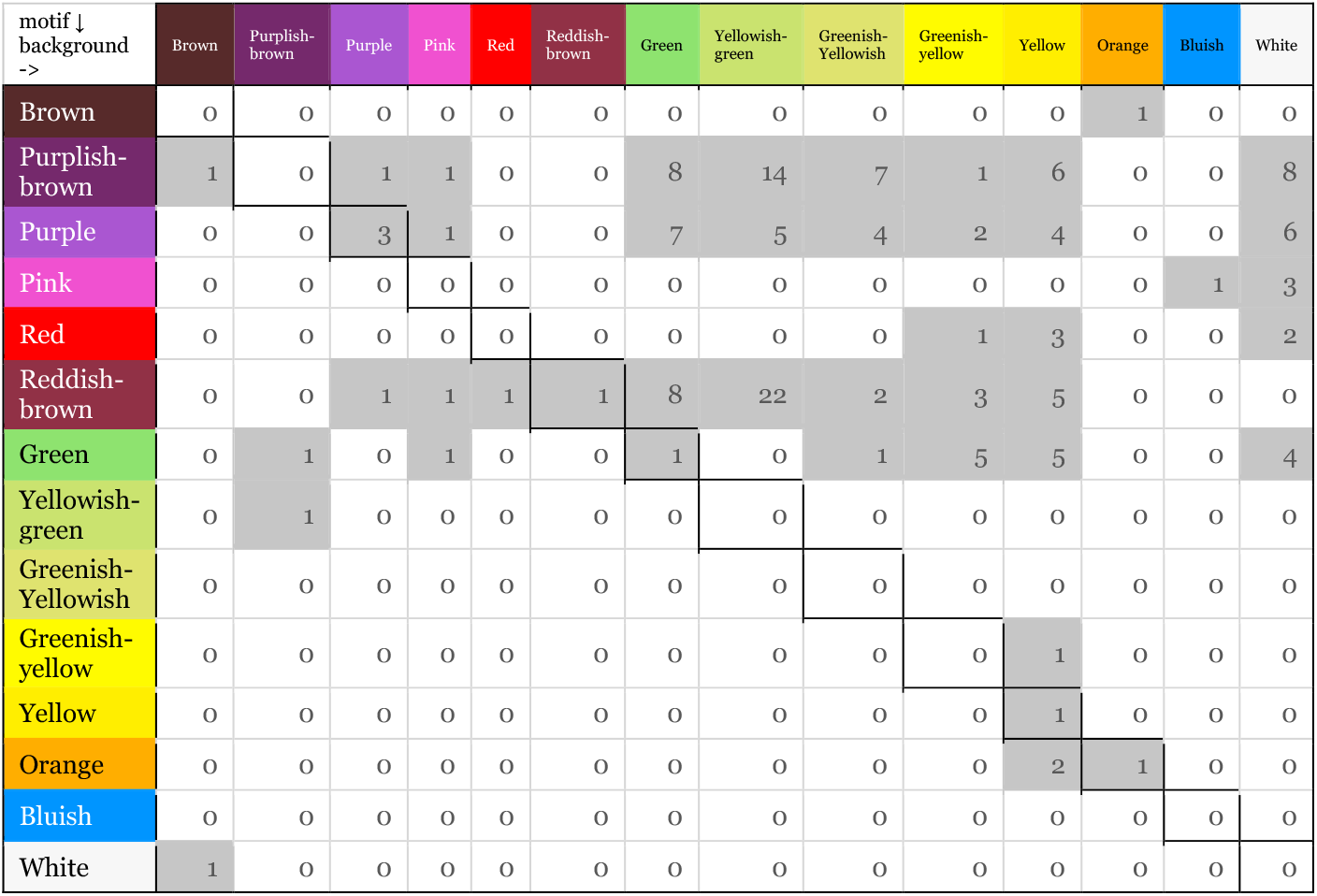
Motif and background color combinations in Fritillaria species. Each number corresponds to the number of recognized species exhibiting this combination.

Furthermore, the diversity of these color patterns is not limited to colors alone, but also extends to shapes, and indeed, the checkerboard pattern is not the only color pattern that exists in *Fritillaria* tepals. We have described seven families of patterns (Fig 2D), which can be grouped into four categories: small periodic patterns (checkerboard, stains, spots), large area patterns (bull’s eye and striped), vascular network coloration (or related to it) and 3D papillae (not color patterns per se, but visible because of their 3D shape and/or the shadows generated by the reliefs).

Interestingly, combinations of several pattern families exist, but some have never been observed, either for adaptive reasons or because they are mechanistically impossible. In particular, the three patterns of small periodic motifs (checkerboard, stained and spotted) are almost never observed simultaneously in an individual, and even if they are, they are never superimposed. This led us to hypothesize that these three families of patterns are produced by the same morphogenetic mechanism, and that the internal or external parameters of the system determine one or the other output.

Finally, the diversity of these color patterns also extends to their location, as shown in Fig 2B. Some species have a visible pattern on both sepals and petals, others only on petals (the reverse has not been observed), some have patterns only on the inner surfaces of tepals, and surprisingly, in at least one species, the pattern is absent from the tepals but present on the peduncle.

The phylogeny of *Fritillaria* mapped with these color and shape characters reveals a probable presence of checkerboard pattern in the common ancestor of the genus (Fig 2C). We also explore color patterns in the *Fritillaria* sister-genus *Lilium*, together with all their sub-family Lilioideae genera. Many periodic patterns are present but no checkerboard pattern was found, suggesting an evolutionary innovation after the separation of *Fritillaria* and *Lilium* lineages:

Tribe: Medeoleae

*Clintonia*: No particular color pattern found

*Medeola*: No particular color pattern found on flowers but reddish bullseyes on bractea in some species like *Medeola virginiana*

Tribe: Lilieae

*Cardiocrinum*: Patterns on the base of the tepals, with spotted and vein patterns.

*Erythronium*: Patterns on the base of the tepals, spotted patterns. Spotted or network patterns on the leaves (e.g., *Erythronium dens-canis*)

*Fritillaria*: (detailed earlier)

*Gagea*: Patterns on the base of the tepals, vein patterns (e.g., *Gagea serotina*) *Lilium*: Spots patterns only above the veins, in particular above vein intersections *Nomocharis*: PGTCPs (Putative-Growth Turing-like Color Patterns), including rosettes (like in *Nomocharis aperta*).

*Notholirion*: Patterns on the base of the tepals with a small spot, sometimes shaped like a circumflex accent (e.g., *Notholirion thomsonianum*)

*Tulipa*: Bullseye patterns

To extract clues on the morphogenetic processes producing *Fritillaria* checkerboard-like patterns, we performed observations on fresh and dissected individuals presenting pattern variability on color and shape (Figure 2.A-C). Figure 2. D (and the corresponding numbered arrows in Figure 2.A-C) summarizes the results in nine pattern observations organized in three axes with their respective deductions and hypotheses: i) organ and tissue location, ii) relationships with vein patterning and iii) geometry characteristics.

Concerning the first axe, we found that in *F. meleagris* the checkerboard pattern (at least its purplish colored part) is borne by the vacuoles of the puzzle-shaped epidermal cells (Fig 3C and 3D), in both upper and lower epidermis (Fig 3C and 3B). Interestingly, upper and lower patterns are superimposed (Fig 3C, right), suggesting a common morphogenesis, either in one or other of these tissues, in both, or even in a third tissue, such as the precursor of the parenchyma which separates the two epidermises. Because the lacunar nature of the spongy parenchyma(15) seems unsuitable for the establishment of a continuous pattern such as the checkerboard pattern of *Fritillaria*, we therefore hypothesize that it is established quite early during tepal development, before the differentiation of the parenchyma. Since the colors are produced late, this also implies a pre-pattern, coded in a way that remains to be determined.

**Fig 3.**
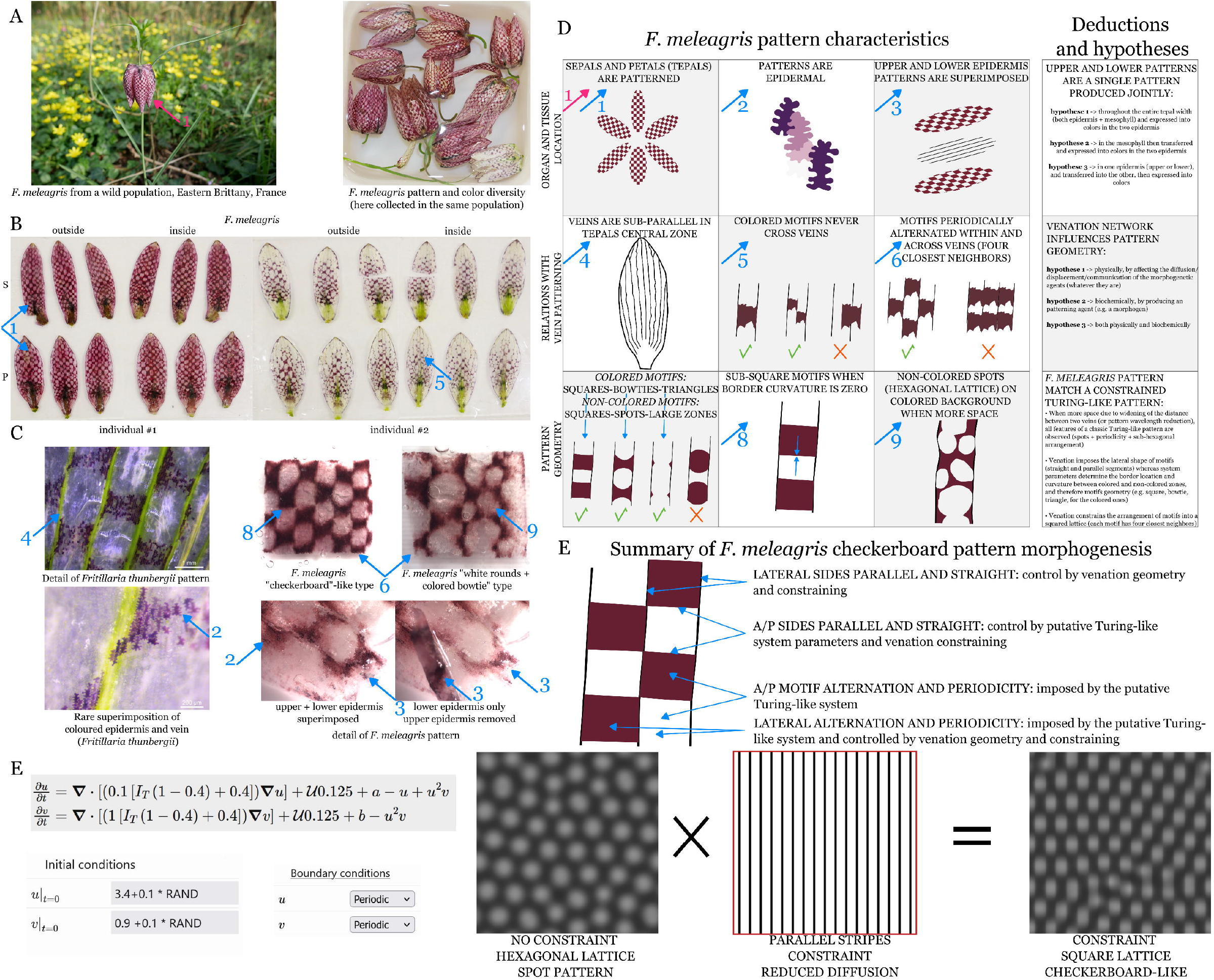
Fritillaria checkerboard patterns. (A) Pictures of *Fritillaria meleagris* from a wild population of Eastern Britanny, France (left) and color diversity from the same population (right) (B) Comparative pictures of sepals (first row) and petals (second row) from two individuals (red morph and white morph), with the respective abaxial/outside and adaxial/inside sides. (C) Magnifications of *Fritillaria thunbergii* (two left pictures) and *Fritillaria meleagris* (four right pictures) color patterns. (D) The nine pattern characteristics with their respective deductions and hypotheses. Numbered blue arrows are dispatched on A-C panels to illustrate the observations. (E) Summary of *Fritillaria meleagris* checkerboard pattern morphogenesis hypotheses (F) Modeling of *Fritillaria* checkerboard pattern by constraining a Turing pattern in a parallel stripes mask. From left to right: system equations, initial and boundary conditions, modeling without constraint, shape of the constraint mask, modeling with the constraint.

Next, we described a clear link between checkerboard and vein patterns: lateral borders of checkerboard squares exclusively coincide with veins, which are sub-parallel in most of the tepals’ surface (Fig 3C and 3D), a characteristic of many monocots (16). It remains then to determine whether this correlation is a sign of chance, of a common inheritance of a third structure that remains currently unknown, or, as we hypothesize, of the influence of one on the other, in this case of the vein pattern on the color pattern. While the vein pattern appears to act as (or to produce) a boundary, it must not be entirely permeable, since the alternation of patterns between two consecutive veins leads us to hypothesize that possible signals can cross (Fig 3C). We have to keep in mind veins are located outside the epidermis, and the latter is continuous so whether they act as a physical, biochemical, or any other type of boundary signal, it will have to be taken into account in the hypotheses. In particular, these observations could acknowledge the hypothesis of a pattern establishment taking place in the parenchyma, as its space is restrained where veins are present.

Finally, while the vein pattern seems to play a direct or indirect role in the final geometry of the checkerboard, it does not seem sufficient to explain all of its characteristics, including the anteroposterior borders of the squares, as well as the alternation of the motifs and their anteroposterior periodicity. To make hypotheses on the characteristics of the morphogenetic process(es) at work, we took advantage of the variations in the geometry of the pattern, whether within the same tepal, or between individuals. We found that the motif geometry of i) the colored areas is a continuum from squares to little triangles via bowties shapes and ii) for the unpigmented areas, from squares to continuous background via truncated disks (Fig 3C and 3D). In some individuals, we observed a very interesting feature in the places where the inter-vein distance is the widest (and/or when the pattern periodic wavelength is smaller): a non-colored spot pattern on a colored background is observed instead. Given the periodicity of this pattern, and the round geometry of the motifs, the variability and the complementarity between colored and non-colored regions, it shares all the characteristics of a Turing-like pattern (17). Indeed, at that time, the latter is the only mechanism demonstrated to produce periodic color dots patterning in Angiosperms (18). Turing pattern relies classically on reaction and diffusion of morphogens (but can be generalized to other scales and natures (19)), and it seems that veins could act as constraining semi-permeable barriers, which therefore shape the pattern into square-like shapes. This notion of constraint in a Turing system has been demonstrated in zebrafish with the role of a lateral structure transforming a spot pattern into stripes (20), and is believed to occur in the lizard *Timon lepidus* to form their scale color pattern (21).

Using Visual PDE (22) and a custom-made Turing system based on Schnakenberg equations, we simulated the effect of a parallel striped constraining on Turing pattern formation (Fig 3E). From an initial classical hexagonal lattice, the constraint transforms it into a square lattice, and spots motifs transformed into rectangle-like motifs, leading to an overall checkerboard-like pattern (Fig 3E)

Fig 3.E them sums up the deductions and hypotheses we made from our observations and simulations.

To produce a final checkerboard-like pattern, several mechanisms need to interact: first, the lateral sides of the squares are produced by the straightness and parallelism of the tepal veins. Secondly, the two other sides are more or less straight, depending on the species and the individual, and might depend on the Turing parameters of the system. Finally, the same Turing-like mechanism seems to be involved to produce an alternation of colored and uncolored zones, characteristics of the checkerboard-like patterns.

### Sarcophaga checkerboard patterns are relying on a structural coloration generated by planar polarity of hair, compatible with the interference between two stripe patterns

Unlike the genus *Fritillaria*, the phenotypic diversity of *Sarcophaga* color patterns is much more limited, both in terms of colors (in this case grey shades) and shapes (23). Furthermore, we observe that many species of other genera of the Sarcophagidae family exhibit patterns that are more or less similar to checkerboards, sometimes without the alternation of dark and light squares, sometimes with spots or triangles instead of squares, or sometimes with only stripes (typically two per segment, aligned from right to left, light in the anterior side and dark in the posterior side). Given this relatively low variability in shapes and colors, we decided against conducting an exhaustive study of the genus and focused instead on a restricted number of species.

Like for *Fritillaria* checkerboards, we perform an in-depth patterns analysis we then summarized in nine observations grouped in 3 categories (1. Macroscopic observations, 2. Links with setae phenotypes and 3. Correlations between colors and setae phenotype) presented in Figure 3.D and in the corresponding arrows (numbered 1 to 9).

First, the checkerboard is located on the cuticle of the dorsal part of the abdomen (Fig 4A and 4D, arrow 1), which is not the only part with a pattern, as the thorax cuticle has stripes, or even intermediate bands, a category of PGTCPs (Putative-Growth Turing-like Color Patterns). We counted 28 squares, divided into three and a half batches of eight squares, across four segments (Fig 4B and 4D, arrow 2). Some sides of the squares match other structural boundaries: the midline and the boundaries between segments. Then, we describe an unexpected feature of *Sarcophaga* checkerboards by observing them under a binocular microscope: some squares could change their colors (or more precisely their shade of grey) depending on the incident light direction (Fig 4A and 4D, arrow 3). Some squares can change from “light” to “dark” through intermediate shade (6 for each octet of squares), while others remain dark regardless of the angle of incidence (the two located in the posterior and lateral corner). In addition, the midline is marked by a thin ever black stripe, as well as two very thin ever black stripes on either side of the midline and parallel to it, located between the first and second columns of squares. Together, this led us to postulate a “physical” coloration (24), as it depends on the angles of illumination and observation, and a morphogenesis that may rely (at least partially) on existing boundaries, such as the midline, and that use the right-left symmetry of the abdomen.

**Fig 4.**
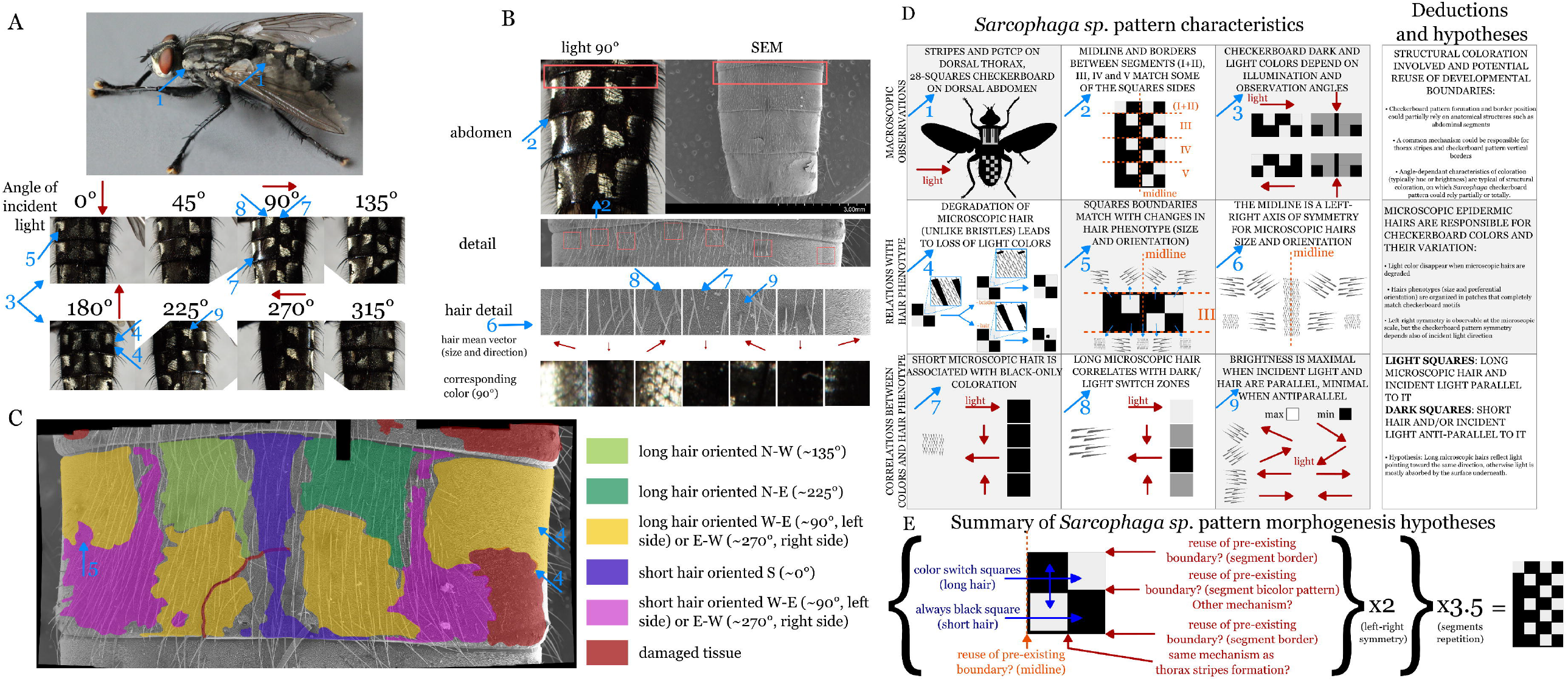
Sarcophaga checkerboard patterns. (A) Pictures of *Sarcophaga sp*. (up) and detail of the abdomen checkerboard pattern from an individual of the MNHN Diptera collection (down). The eight pictures correspond to the same abdomen taken with different incident light angles. (B) Correspondence between color and setae hair phenotypes, under a 90 degrees angle of light illumination. Up left picture and fourth row: light microscopy, all the other pictures: SEM. (C) Assembly of SEM pictures and superimposition of colors corresponding to the phenotype of the setae of the 2^nd^ abdomen segment. (D) The nine pattern characteristics with their respective deductions and hypotheses. Numbered blue arrows are dispatched on A-C panels to illustrate the observations. (E) Summary of *Fritillaria meleagris* checkerboard pattern morphogenesis hypotheses

To further explore the origin of this changing pattern composed of grey shades, we took a microscopic look by examining samples of *Sarcophaga* sp. abdomens under a scanning electron microscope. In particular, we explored the characteristics of the small epidermal hairs covering much of the external surface of the abdomen, called setae, which are then strongly suspected of contributing to the checkerboard pattern.

First, by examining partially damaged museum specimens, we were able to demonstrate that areas where the setae are damaged lose their ability to change their shade and remain constantly dark (Fig 4A and 4D, arrow 4). In contrast, when the large “bristles” (another type of epidermal hair) are damaged, this has no visible effect on the pattern or the coloration. Next, we observed that the setae were organized into large areas characterized by size (large and overlapping, or small and revealing the surface underneath) and preferred direction (also known as planar polarity (25)), and that the boundaries between areas corresponded perfectly to the edges of the checkerboard squares (Fig 4A, 4C and 4D, arrow 5). Finally, we observed a right-left symmetry in these microscopic characteristics, with long setae oriented N-W (∼135°), N-E (∼225°), W-E (∼90°) or E-W (∼270°) (Fig 4B and 4D, arrow 6). We can note that the areas closest in terms of direction are located diagonally to each other, which surely plays a role in the alternation of dark and light squares in the checkerboard pattern. There is therefore a perfect correlation between microscopic boundaries based on the setae phenotypes and macroscopic boundaries based on color. To provide clues as to the existence of a causality between micro and macro scales, we explored in detail the microscopic characteristics of setae in relation to the direction of light.

We first observed that the areas that remain dark regardless of the incident light conditions (i.e., the midline, the two parallel lines, and the lateral-posterior squares) always have short setae that do not cover the entire surface, revealing the hairless parts of the epidermis (Fig 4A, 4B and 4D, arrow 7). On the contrary, the squares which change of shade depending on the light are composed of long, lying setae partially overlapping and hiding the hairless part of the epidermis. When incident light rotates through 360 degrees, each zone passes once through a maximum brightness leading to a light square and once through a minimum brightness leading to a dark square, with shades of gray in between (Fig 4A, 4B and 4D, arrow 8). Finally, squares are at maximum brightness when the light points in the same direction, and at minimum brightness when the light is anti-parallel (Fig 4A, 4B and 4D, arrow 9). All these observations lead us to propose the following hypotheses:

The hairless part of the epidermis absorbs light (for example, via melanin pigmentation), while the hairs reflect incident light (without any particular wavelength preference, hence the shades of gray). When light is parallel to the direction of the hairs, it is reflected on their surface and reaches the observer. Conversely, when it is anti-parallel to the hairs, it penetrates inside and beneath the setae network and is absorbed or prevented from escaping, leading to a lack of light to the observer.

Fig 4E sums up the deductions and hypotheses we made from our observations.

First, we simplify the pattern as the repetition of a unit of four independent squares, then doubled using left-right symmetry, then multiplied by 3.5 using the abdomen segmentation, leading finally to 28 squares. The outer boundaries of the 4-square unit all coincide with an existing structural boundary (midline, boundaries between segments and dorso-ventral boundary), so we hypothesize that they are used for the pattern. On the contrary, the two internal perpendicular boundaries do not coincide with any observable structural frontier. However, many species possess lateral stripes on the abdomen(26,27), therefore the mechanism that produces them could be reused. Concerning the anterior-posterior boundaries, we hypothesize that the mechanism that produces the thorax(28) ones could be co-opted here in the abdomen. Finally, the combination of these perpendicular stripes could code for the length and orientation of the setae, which by extension codes for their behavior with respect to the angle of incident light their and therefore for the resulted color (or shade).

## Discussion

### Comparing the morphogenesis of Fritillaria and Sarcophaga checkerboards in a standardized flowchart called FORMula

Through this study, it became clear that the morphological similarity of the checkerboards of the two genera studied was not reflected at the level of morphogenetic processes, where evolutionary convergence appears to be at work. We defend the idea that a finite number of morphogenetic processes are responsible for the forms and patterns of the living world, and that these can be defined and classified into universal categories (29,11). Furthermore, by summarizing the morphogenesis of a form or a pattern into different sequential stages, we can synthesize it using a flowchart, which we propose to call the FORMula (Fig 5A and 5B). We defined two types of morphogenetic processes, generator and modulator (the first one being sufficient to produce a shape contrary to the second one), respectively represented with a pink trapeze and a blue rectangle with brackets. Shapes (which include both forms and patterns, the latter being characterized by two or more internal states possible, typically colors) are represented by a yellow rectangle. Morphogenetic processes are able to generate shapes from scratch (represented by ‘Ø’ symbol) or to modify existing shapes. Both are represented by an arrow. Two or more processes can interact (represented by ‘X’ symbol) or add (represented with a ‘+’ symbol) to form the resulting shape. Morphogenetic processes, in particular modulators, can also have its own shape, which is represented by a segment which precede them (e.g. putative reduced diffusion in *Fritillaria* possess the shape of the vein sub-parallel network). For the final shape of interest, here the checkerboard pattern, we detail the three necessary and sufficient characteristics, to be able to precise the respective roles of morphogenetic processes. For example, in *Fritillaria*, we hypothesize that the reduced diffusion caused by the vein network is not responsible for color alternation, but for both the square lattice and the square shapes of the motifs. Putative Turing-like process participates to the establishment of all of the three characteristics. These flowcharts can be transcribed into text form (e.g. for *Fritillaria* checkerboards in Fig 5C), while containing the same amount of essential information (descriptive names of processes and illustrations are excluded for clarity reasons). Theoretically, the entire morphological development of an individual could be represented in this form. By analogy with phylogenetic trees and cell lineage trees, an appropriate name might be “morphogenetic tree.” Of course, this representation is an intellectual construct of processes that are more complex than the information contained in the FORMula and, moreover, until functional studies provide direct evidence, many areas of the diagram remain at the hypothesis stage. Nevertheless, it allows knowledge and hypotheses to be synthesized while clearly differentiating between them, and facilitates comparative analyses between species. Finally, like in this study, FORMula allows to pinpoint the links between different shapes to be directly represented. We hypothesize that many biological structures, once formed, serve by their own shape as constraints, bases, boundaries, and/or signals for the morphogenesis of other biological structures. This intertwining of shapes is understudied in morphogenesis, particularly due to the obvious focus on ‘generator’ morphogenetic processes in the first instance, whereas what we have referred to above are more like ‘modulators’ of shapes. By analogy with phylogenetic trees, numerous ‘horizontal transfers’ of shapes appear to occur during the morphogenesis of organisms—at least, that is our working hypothesis—and we encourage the community to explore these links in an attempt to assess their significance.

**Fig 5.**
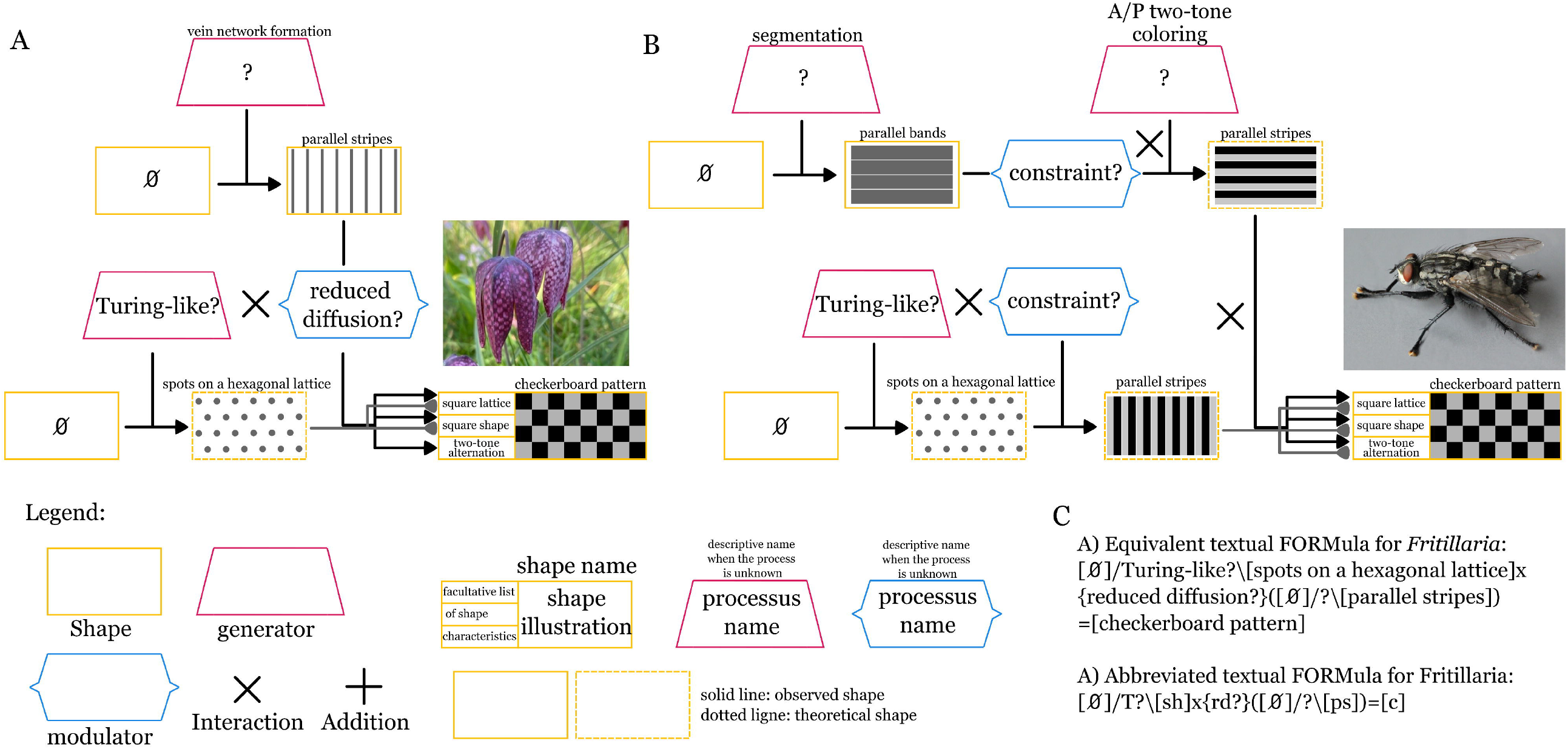
FORMula, a standardized and logical representation of morphogenesis. Yellow rectangle: shape, pink trapeze and blue bracket-shaped rectangle: morphogenetic processes (respectively generator and modulator). (A) FORMula of *Fritillaria meleagris* checkerboard pattern (B) FORMula of *Sarcophaga sp*. checkerboard pattern (C) Equivalent textual FORMula for *Fritillaria meleagris*

## Materials and methods

### Scanning Electron Microscopy

A total of three abdomens were sampled from *Sarcophaga* sp. museum specimens (National Museum of Natural History, Paris, France). Samples were mounted on the aluminum stubs with colloidal graphite, sputter-coated with platinum using an EM ACE600 fine coater (Leica, Wetzlar, Germany), and observed using a SU3500 scanning electron microscope (SEM, Hitachi, Tokyo, Japan).

### Light Microscopy and photography

*Fritillaria* thunbergii: a total of 20 tepals were sampled from 8 specimens of *Fritillaria* thunbergii, harvested in the botanical gardens of the University of Tokyo (Koishikawa Botanical Gardens, Tokyo, Japan). Tepals gas were extracted by a vacuum pump to make the tissues more transparent and suitable for observation. Abaxial and adaxial sides were mounted in water and observed under light microscopy.

*Fritillaria meleagris*, herbarium specimens: A total of 8 tepals were sampled from 3 specimens of *Fritillaria meleagris*, from museum specimens (National Museum of Natural History, Paris, France), after a survey of hundreds of museum specimens. They were rehydrated and observed under light microscopy.

*Fritillaria meleagris*, fresh specimens: A total of 31 specimens were harvested from a wild population in Eastern Britanny, France. They were chosen to be representative of all the developmental stages of tepal morphogenesis and to present as much color and shape variability as possible. All tepals (8 per flower, one flower per specimen) were mounted in water between two glass sheets and photographed on both abaxial and adaxial sides with scales. 16 sepals and 16 petals were selected to be mounted in distilled water between a slide and a cover slip. They were observed under a binocular microscope. To observe independently abaxial and adaxial epidermis of the same tepal, we separate them with the sharp edge of a glass cover slip.

### Biodiversity surveys among Eukaryotes and Fritillaria genus

In order to take as comprehensive an approach as possible, we used multiple sources: first, scientific literature, using keywords such as “checkerboard,” “checkered,” “check,” “square,” and “*Fritillaria*” with the word “pattern,” then adding the names of clades in Latin or English, such as ‘insecta’ or “insect.” We then consulted several experts from different taxonomic groups, particularly those working at the National Museum of Natural History of Paris, France. Finally, we use secondary sources like online encyclopedias, Wikipedia (https://en.wikipedia.org/) and Wikimedia Commons, and image browsers. For each candidate species, we double checked with scientific sources. In addition, some groups had already been explored for another study focused on other patterns, PGTCPs, but since the author had already begun this study on checkerboards, he systematically noted those he found. Thus, the 6,500 species of mammals were explored, as were the 30,000 species of orchids, and all families of angiosperms and orders of animals were explored using keywords. The fact that we found checkerboard patterns in small numbers but in groups that are highly representative of the macroscopic biodiversity of angiosperms and animals reinforces our belief that we have not overlooked too large a proportion of the clades exhibiting these patterns.

Phylogenetic relationships between species were built with phyloT (V2, https://phylot.biobyte.de/) based on NCBI taxonomy. Phylogenetic trees were plotted and annotated in iTOL (Interactive Tree Of Life, version 6.8.1, https://itol.embl.de/)(30)

### Modeling

All the model parameters are available at: https://visualpde.com/sim/?mini=lR1qNaKv

We use the software Visual PDE (22) and a custom-made Turing system based on Schnakenberg equations, and we simulated the effect of a parallel striped constraining on Turing pattern formation thanks to a mask containing 16 black stripes on a white background designed on GIMP (version 3.0.4). Each black pixel corresponds to a reduction of 60% of the diffusion rate compared to the control, whereas white pixels kept 100% of the diffusion rate.

## Acknowledgments

We deeply thank Florian Jabbour for bringing *Fritillaria* patterns to our attention and for his advice for Scanning Electron Microscopy.

We deeply thank Thierry Deroin for the help in the rehydrating and preparation of museum specimens

We deeply thank the National Museum of Natural History, Paris, France for the access to the collection and the authorization for collecting museum specimens for destructive experiments

We deeply thank Pr. Hirokazu Tsukaya, the University of Tokyo and the Koishikawa Botanical Gardens of the University of Tokyo for the authorization for collecting specimens of *Fritillaria thunbergii* specimens

We deeply thank Nicolas Bekkouche and Paul Chatelain for their suggestions for the species survey

## Supporting Information

### S1 Table Fritillaria color patterns survey

The 164 recognized species of *Fritillaria* have been screened for 1) Distribution area 2) Presence of a particular color pattern: Yes checkerboard (Y), Other (O), None (N). 3) Organ and side where a pattern is visible: Sepal adaxial. (SE), Sepal abaxial (SI), Petal adaxial (PE), Petal abaxial (PI), All Organ and side where a pattern is visible. 4) Other organs bearing checkerboard, check (or tessellated) patterns, 5) Geometrical description, 6) Pattern variability (estimation among individuals), 7) Motif color, 8) Background color and 9) Color and literature references.

